# Nightingales imitate the duration of whistle syllables

**DOI:** 10.1101/2025.04.12.648498

**Authors:** J.S. Calderon-Garcia, G. Costalunga, T.P. Vogels, D. Vallentin

## Abstract

While real-time vocal adjustments are crucial for interactive communication, not much is known about the spectral and temporal vocal flexibility of animals. Here, using a combination of field song recordings and controlled playback experiments, we show that wild nightingales imitate ‘whistle song’ syllable durations in real time. However, when exposed to playbacks with variations in whistle pitch and duration that are beyond their natural range of vocalizations, nightingales emulated temporal or spectral features, approximating the whistle playbacks within the constraints of their natural whistle song repertoire. Our findings reveal a previously unknown dimension of real-time vocal flexibility in songbirds and suggest complex auditory-motor integration during song interactions.

## Main

Real-time vocal modifications of acoustic features play a key role during vocal interactions^1^. Several animals including humans are known to precisely time their vocalizations during vocal exchanges^2,3^. However, little is known about the ability of songbirds to spontaneously adapt the temporal structure of their vocalizations in response to auditory stimuli^4^. We explored this phenomenon in wild nightingales, a species known for natural song-matching, during which they listen to and immediately replicate the songs of other nightingales^4^. We focused on their “whistle songs”, sequences of unmodulated tonal syllables (Fig 1A), that nightingales are known to pitch-match in real time during counter-singing duels^5^. To investigate if nightingales can also adjust whistle syllable duration, we recorded vocal responses of wild nightingales to artificial whistle playbacks during their breeding season in Germany. In the absence of stimuli, we observed three clusters of whistle syllable durations: short syllables (shorter than 150 ms), medium syllables (between 150 and 300 ms), and long syllables (longer than 300 ms) (Fig 1A, Fig S1A, B, see methods). To test whether nightingales can shift their natural distribution of whistle syllable durations, we presented a battery of playbacks consisting of whistle syllables with durations corresponding to the four less frequent temporal regimes (Very-Short, Medium-Short, Medium-Long, Very-Long) (Fig 1B, C, Fig S1E). Although the overall range of whistle syllable durations in response was broad, we observed a notable shift in the distribution of duration of the whistle syllable responses towards the duration of the Very-Short, Medium-Short and Very-Long playbacks (Fig 1D, Fig S2A-C). We developed an ‘occurrence index’ to assess if whistles of specific durations occurred more or less frequently during playback conditions than expected by chance; an occurrence index below 1 indicated fewer, equal to 1 indicated the same, or above 1 indicated more whistles in the experimental group compared to the natural whistle distribution (see methods). The observed occurrence index greater than one in the vicinity of playback durations indicated a targeted shift of whistle syllable duration toward the stimuli duration (Fig 1D), demonstrating that nightingales can flexibly mimic temporal aspects of whistle songs in response to whistle stimuli. This capacity for "one-shot" duration imitation has not been previously observed in animals without explicit training.

**Figure 1.**
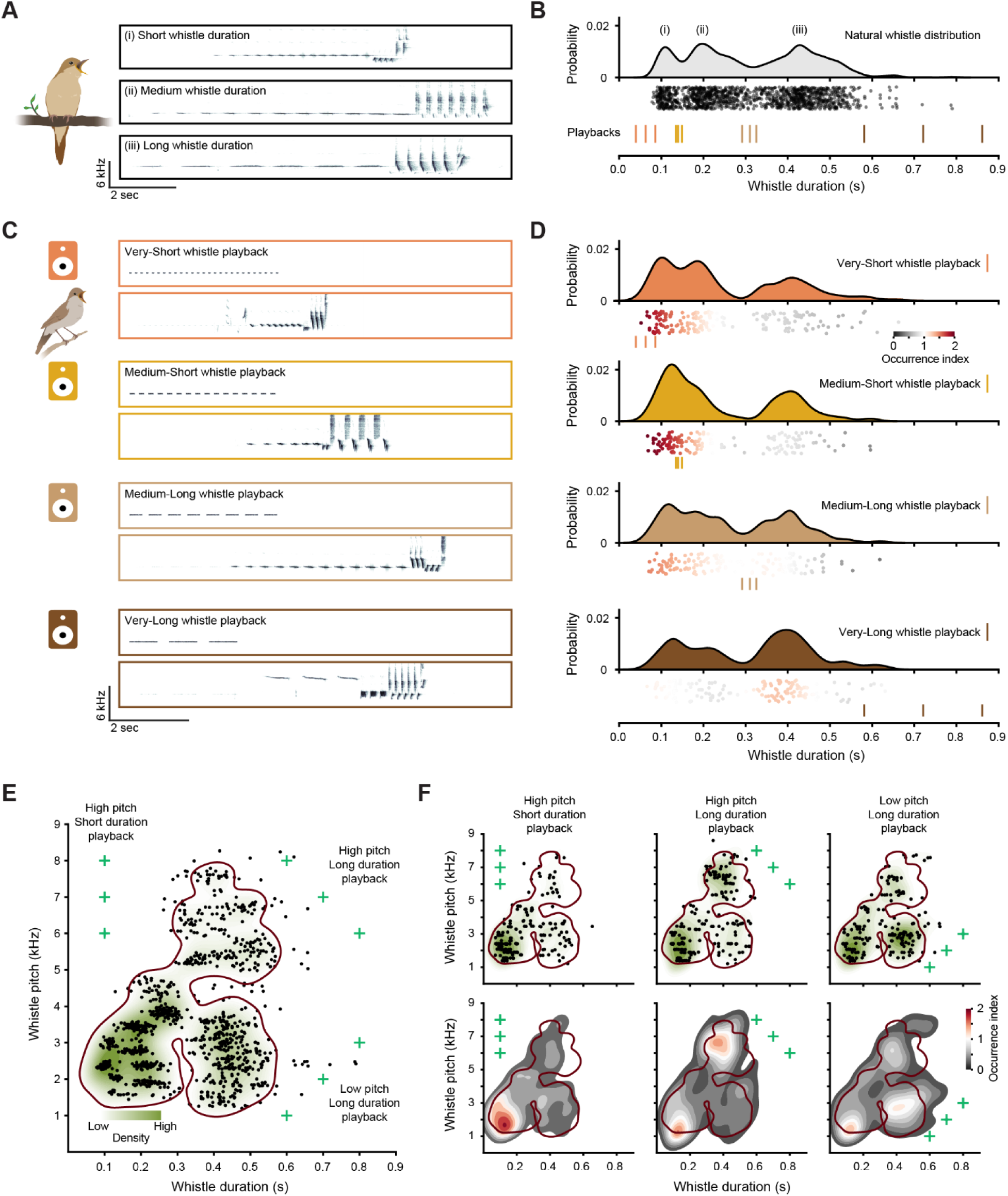
Flexible imitation of whistle syllable duration in nightingales. (A) Three examples of whistle song spectrograms with (i) short <150 ms), (ii) medium (150-300 ms), and (iii) long (>300 ms) whistle syllable durations. (B) Top: Natural distribution of whistle syllable durations (0.294±0.124 s; n=1237 from N=11 birds). The three modes of the probability distribution correspond to the short (i), medium (ii) and long (iii) syllable durations shown in (A). Middle: Each black circle corresponds to the median whistle syllable duration of whistle syllables of individual whistle songs. Bottom: Vertical colour lines represent the durations of playback stimuli (Orange=Very-Short; Gold=Medium-Short; Sand=Medium-Long; Brown=Very-Long playbacks). (C) Example spectrograms of playbacks and nightingale responses for the four different playback regimes. (D) Probability distributions of nightingale whistle responses to each playback regime: Very-Short (0.194±0.096 s, n=172 from N=7 birds); Medium-Short (0.185±0.075 s, n=155, N=7); Medium-Long (0.236±0.112 s, n=153, N=7); Very-Long (0.346±0.111 s, n=152, N=7). Individual whistle song responses were color-coded based on the relative frequency of occurrences within the song duration compared to the naturally observed distribution, calculated as the occurrence index (See methods). (E) Natural distribution of whistle syllable duration and pitch (n= 1237, N=11). Each black circle corresponds to the median whistle syllable duration and pitch of whistle syllables in individual whistle songs. The bright green crosses represent the duration and pitch of the three types of playback stimuli (High-pitch/Short-duration, High-pitch/Long-duration, Low-pitch/Long-duration). The green shading represents the Kernel Density Estimation (KDE) of the whistle distributions in the soundspace (See methods). The brown line encloses 90% of the control KDE density. (F) Top: Distribution of whistle syllable duration and pitch in response to the three types of playback stimuli (High-pitch/Short-duration, n=193, N=8; High-pitch/Long-duration, n=194, N=8; Low-pitch/Long-duration n=190, N=8) represented as in (E). Bottom: Density estimation of responses color-coded based on the relative density of occurrences compared to the naturally observed distribution, calculated as the occurrence index of observed to expected densities (See methods). The brown line encloses 90% of the control KDE density.

So far all our playback stimuli consisted of the same pitch, 2390 Hz, that nightingales can readily match^5^. However, whistle songs naturally produced by singing nightingales comprise a broad and multimodal distribution of pitches and durations (Fig 1E). To assess this multimodality, we applied k-means clustering to the distribution of whistles syllables in the soundspace (defined here as the duration-pitch feature space). We found three main clusters (C1=Low pitches/Short duration syllables; C2=High pitches syllables; C3=Low pitches/Long duration syllables, Fig S1F, G). To explore a potential trade-off between pitch-matching and temporal imitation, we presented birds with a set of three playback types from three areas in the soundspace (High-pitch/Short-duration, High-pitch/Long-duration, Low-pitch/Long-duration), combining common and uncommon temporal and spectral features (Fig 1E, Fig S2D). In this scenario, nightingales could modify their whistle responses in syllable duration, pitch, neither, or both. A trade-off between spectral and temporal imitation would reveal how nightingales prioritize adjustments to stimuli out of their natural production range.

We observed a shift in the soundspace probability density distribution of whistles in response to the three playback types compared to the natural whistle distribution (Fig 1F, Fig S2E). For each playback type, we computed the proportion of responses belonging to each of the three clusters defined by the k-means algorithm and compared them to the expected proportions from shuffled datasets (Fig S2F, See methods). When exposed to High-pitch/Short-duration whistle playbacks (Fig 1E), nightingales targeted the duration but not the pitch (Fig 1F, first), resulting in an increase in whistle responses in the C1 cluster and a reduction of whistle syllables in the C2 cluster (Fig S2F). Conversely, for High-pitch/Long-duration playbacks (Fig 1E), the birds matched the pitch but not duration (Fig 1F, second), increasing the production of whistles in the C2 cluster (Fig S2F). Finally, for Low-pitch/Long-duration playbacks (Fig 1E), nightingales matched the pitch and lengthened whistles, but not to the full syllable durations featured in the playback (Fig 1F, third), demonstrated by a reduction of whistles in the C1 and C2 clusters, and an increase of whistles in cluster C3 (Fig S2F). These findings show that, when presented with artificially constructed whistle songs comprising acoustic features outside of their natural whistle syllable repertoire, nightingales can adapt their responses by projecting them back into their natural whistle soundspace through selective imitation of temporal and/or spectral features. Our results suggest a flexible but potentially constrained strategy for prioritizing acoustic features to emulate sounds outside of the natural vocal production range of nightingales.

We concluded that counter-singing nightingales exhibit considerable vocal complexity by not only pitch-matching^5^, but also through adjusting the duration of syllables in their whistle songs. The ability to specifically imitate both the temporal or spectral features of whistle stimuli suggests a capacity for auditory discrimination, allowing nightingales to isolate individual vocal features such as pitch and syllable duration from complex sounds in real time--as much as they are physically able--instead of merely mimicking overall patterns.

Our current understanding of the neural circuit underlying song production is largely based on songbird species with limited flexibility in singing behaviour, such as zebra finches^6,7^. In these finches, the temporal structure of vocal motor output is tightly controlled by the premotor nucleus HVC (proper name)^8^, and minor frequency or temporal changes in song structure can only be induced through long-term reinforcement learning^9,10^. This contrasts with the real-time flexibility observed in nightingales, suggesting rapid mechanisms for adjusting vocalizations in response to auditory input. Our results from counter-singing in nightingales indicate a species-specific adaptation in neural motor control for vocal flexibility that enable real-time production of songs to imitate temporal and pitch variations in ambient whistle songs. Investigating such behaviour and its neural correlates could shed light on broader mechanisms for song production, vocal learning and real-time acoustic modifications during interactive communication.

### METHODS

## Experimental animals and ethics

We studied a wild population of Common nightingales (*Luscinia megarhynchos*) in the southwestern part of Brandenburg, Germany between April and May in 2020 and 2024. All recordings and playbacks were performed in accordance with the local authorities (Landesamt für Umwelt - Land Brandenburg LFU-N4 4730/14+5#181132/2021).

## Generation of artificial playbacks

### Artificial whistles generation

Artificial whistle syllables were synthesized using Python’s NumPy and SciPy libraries. A white noise signal was generated at a 44.1 kHz sampling rate. This signal was then bandpass filtered with a 120 Hz bandwidth centred on the target pitch. A 0.01 s Hann window was applied for boundary fading. Finally, the resulting signal was amplitude-normalized. We stablish the silent interval duration based on the relationship between whistle syllable duration and the duration of the subsequent silent interval in the control data. (Fig. S1E). Control data exhibited a positive correlation (R² = 0.43), indicating that shorter/longer whistle durations were associated with correspondingly shorter/longer silent intervals, following the linear relation:

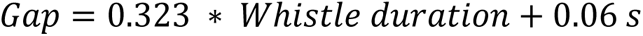

For both experimental paradigms, the inter-syllable silent interval within each playback was parametrically defined as a function of the respective whistle syllable duration. Subsequently, these silent intervals were appended to their corresponding whistle syllables, and the total number of whistle-interval units was given by the maximum number of entire units that fitted within 4 s.

### Playback stimuli for experiment 1

Twelve playback stimuli were generated, each with distinct whistle durations and a constant frequency of 2390 Hz. Whistle durations were selected based on the trimodal distribution observed in the control dataset. According to cluster analysis of control whistle syllable durations (see: Clustering of whistle syllable durations (Experiment 1)), three duration categories were defined: short (<0.14 s), medium (0.14-0.31 s), and long (>0.31 s). Playback durations were then determined by sampling within each category, specifically including the minimum and maximum values, the 5th and 95th percentiles, and interpolated midpoints between selected values. This resulted in four distinct duration ranges for playback whistle syllables: Very-Short (0.04 s, 0.06 s, 0.08 s), Medium-Short (0.13 s, 0.14 s, 0.15 s), Medium-Long (0.30 s, 0.31 s, 0.32 s), and Very-Long (0.58 s, 0.72 s, 0.86 s).

### Playback stimuli for experiment 2

Stimuli were designed with durations and/or pitches outside the natural soundspace of nightingale whistle songs. These regions were defined as areas where whistle syllable density fell below 10%. Specifically: for High-Pitch, Short-Duration three playbacks were created with a fixed whistle duration of 0.14 s and varying frequencies: 6 kHz, 7 kHz, and 8 kHz; for High-Pitch, Long-Duration three playbacks were generated with durations of 0.6 s, 0.7 s, and 0.8 s, and corresponding frequencies of 6 kHz, 7 kHz, and 8 kHz; for Low-Pitch, Long-Duration three playbacks were produced with durations of 0.6 s, 0.7 s, and 0.8 s, and corresponding frequencies of 1 kHz, 2 kHz, and 3 kHz.

## Audio recordings

Audio recordings (16-bit precision at 44.1 kHz sampling rate) from a total of 22 wild male nightingales were obtained during the mating season (April/May) of 2020 and 2024. Recording sessions were conducted between 11pm and 4am CET+1. Each nightingale was recorded with a directional parabolic microphone equipped with a windshield (Stereo MK3, Telinga, Sweden), connected to a battery-driven pre-amplifier (Roland Duo-Capture EX, Roland, Japan) and a laptop computer. Signals were acquired using Audacity v.2.4.2 (https://www.audacityteam.org/) and encoded as stereo .wav files (one channel for the playback and one for the bird).

## Control recordings

Recordings from 11 naturally singing nightingales, acquired in 2020, were used as control data.

## Song playback experiment

Playback experiments with 11 birds were performed as previously described. In brief, a speaker (JBL, Harman International Industries, USA) was connected to the pre-amplifier and placed at ∼10 meters from the singing bird. The playbacks were manually triggered depending on the singing behaviour of the nightingales, to avoid overlaps with the birds’ songs. Each bird received 20 repetitions of every playback stimulus. Experiment 1 consisted of 12 different stimuli that were randomly shuffled across the execution of the experiment for a total of 240 playback stimulations. Experiment 2 consisted of 9 different stimuli.

## Audio data processing and analysis

Audio recordings were processed and analyzed using Audacity, Avisoft SASLab Pro 5.2 (R. Specht, Berlin, Germany) and Python.

As previously described^5^, stereo files were divided in mono files and high pass filtered (frequency=1000 Hz, roll-off=6 dB, High-Pass Filter build-in function of Audacity) and noise reduced to remove environmental noise (noise reduction=12dB, sensitivity=6.00, frequency smoothing=3, noise reduction build-in function of Audacity). Sonograms were examined in Audacity for visual scoring of whistle responses. A song was considered a whistle response to playbacks when the following criteria were met (1) being a whistle song (containing frequency unmodulated whistle syllables) and (2) being an immediate response to the playback (i.e. the first or second song sung by the bird after the termination of the playback). Onset and offset of these songs were visually inspected and manually annotated. Individual whistle syllables were manually segmented from whistle songs in Audacity. Avisoft was used for automatically measuring the average frequency and duration of individual syllables.

## Quantification and statistical analysis

Data analysis and statistical computations were performed using Python. All values are reported as median ± median absolute deviation, if not noted otherwise.

### Clustering of whistle syllables duration (Experiment 1)

A Gaussian Mixture Model (GMM) was applied to analyze the distribution of whistle syllable durations in naturally singing nightingales, utilizing the Gaussian Mixture function from the scikit-learn (sklearn) Python library. GMMs were fitted with varying numbers of components (1 to 9), and the Bayesian Information Criterion (BIC) was calculated for each model. The optimal number of components, k, was determined as the value that minimized the BIC score. This analysis yielded k = 3, with component boundaries at 0.14 s and 0.31 s (Fig. S1A). These boundaries facilitated the categorization of natural whistle durations into three distinct groups: short (<0.14 s), medium (0.14-0.31 s), and long (>0.31 s). Analysis of individual nightingales also revealed a trimodal distribution of whistle durations, consistent with the population data (Fig. S1B)

### Clustering of whistle syllables duration and frequency (Experiment 2)

To analyse the distribution of whistle syllables within the soundspace, k-means clustering was employed (Fig. S1F). The optimal number of clusters (k) was determined using silhouette analysis, which assesses cluster consistency. The mean silhouette score was calculated for cluster counts ranging from 2 to 9, and the cluster count yielding the highest mean score was selected as optimal (k=3). Cluster boundaries (C1, C2, and C3) were defined by Voronoi regions, which encompass all points closer to a given centroid than to any other within the soundspace. Due to the Euclidean distance metric used by the k-means algorithm, these cluster boundaries are linear segments. Analysis of individual, naturally singing nightingales confirmed the preservation of the previously described whistle syllable distribution within the defined soundspace. (Fig S1G).

### Whistle syllable duration variation within whistle songs

Whistle syllable durations within a song exhibited some variability. When examining the duration difference between consecutive whistle syllables, a gradual transition was observed, with a mean difference of 0.008 s and a standard deviation of 0.044 s (Fig. S1C). The duration difference between the first and last whistle syllables in each song revealed a mean duration increase of -0.047 s with a standard deviation of 0.1 s (Fig. S1D). Consequently, the median syllable duration of each song was chosen for subsequent analysis. This metric preserved the previously identified short-medium-long clustered distribution (Fig. 1B), provided a representative measure of individual songs, and ensured inter-song comparability.

### Generating shuffled data

In the playback experiment, the median durations and pitches of observed whistle response syllables were randomly sampled with replacement from the responses dataset and assigned to the corresponding playback durations and pitches. The number of whistle match occurrences was preserved for each playback pitch. To extract the respective parameters the shuffling was repeated 10000 times.

### Occurrence index of durations (Experiment 1)

A comparative density analysis of median whistle durations between control and experimental conditions was conducted using Kernel Density Estimation (KDE) and subsequent KDE ratio calculations. KDE distributions were generated for whistle responses to playback stimuli, stratified by duration category. These experimental KDE distributions were then compared to a control KDE, derived from control data, by calculating a pointwise KDE ratio. The occurrence index is defined as the pointwise division of the experimental KDE (*KDE exp (duration)*) by the control KDE (*KDE control (duration)*):

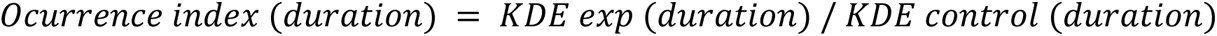

This index captured relative density changes across playback conditions and informed how the experimental distributions for each playback category deviated from the control. An index below 1 indicates fewer whistles in the experimental group, an index equal to 1 indicates no difference, and above 1 indicates more whistles in the experimental group compared to the control.

This approach was employed for a detailed comparative analysis of distributional shifts, specifically highlighting regions of increased or decreased density relative to the control distribution. For visualization purposes, the occurrence index values were clipped to the range of 0 to 2.

### Occurrence index of whistle durations and pitch (Experiment 2)

For each playback region (High-Pitch/Short-Duration, High-Pitch/Long-Duration, and Low-Pitch/Long-Duration) the joint probability density function of whistle duration and pitch was estimated using a Gaussian Kernel Density Estimation (KDE). This KDE was computed over a grid spanning the soundspace, with duration values ranging from 0 s to 0.9 s and pitch values ranging from 0 kHz to 9 kHz. This yielded the estimated experimental probability density, Z exp (duration, pitch). An analogous analysis was performed on the control data to obtain the control probability density, *Z control (duration, pitch)*. at each grid point. Subsequently, an occurrence index was calculated as follows

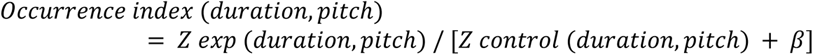

The small constant β=0.0003 prevents division by zero. The occurrence index was clipped within 0 and 2. This index provides insights into how the distribution of experimental responses differs from the control. An index greater than 1 indicates areas where whistle responses are more concentrated for each of the different playback regions. Conversely, an index below 1 highlights region where responses are less frequent. When the index is close to 1, it suggests no significant difference between the experimental and control distributions.

### Statistical analysis for Experiment 1

The statistical significance of observed median whistle durations across temporal categories (Very-Short, Medium-Short, Medium-Long, and Very-Long) was assessed. A null distribution was generated through 10,000 bootstrap resamples of the data (see: Generating shuffled data), and the median was calculated for each category within each resampled dataset (example shuffled distribution in Fig. S2B). P-values were then determined by comparing the observed median for each category to the null distribution

of medians derived from the resampled data (Fig. S2C). For the Very-Short category, a left-tailed hypothesis test was employed, with the p-value representing the proportion of resampled medians less than or equal to the observed median. For the Very-Long category, a right-tailed test is used, where the p-value is the proportion of shuffled medians that are greater than or equal to the observed median. For the other two categories, Short-Med and Med-Long, a two-tailed test is applied, which accounts for deviations in both directions.

### Statistical analysis for Experiment 2

The statistical significance of whistle responses to playback stimuli outside the nightingale’s natural soundspace (High-Pitch/Short-Duration, High-Pitch/Long-Duration, and Low-Pitch/Long-Duration) was evaluated. For each playback region, whistle responses were classified into the three previously defined soundspace clusters (see: Clustering of whistle syllable duration and frequency (Experiment 2)). The percentage of whistle responses within each cluster was calculated for each playback region. Subsequently, the data were subjected to 10,000 shuffle iterations (see: Generating shuffled data), and the percentage of whistle responses within each cluster was recalculated for each shuffled dataset (example shuffled distribution in Fig. S2E). This process generated a null distribution of expected whistle response percentages for each cluster. The p-value in this analysis was computed using a two-sided test to determine whether the observed cluster proportions for each playback region significantly deviated from the null distribution (Fig S2F).

## CODE AND DATA AVAILABILITY

Code and data for this project are available at https://github.com/vallentinlab

## ACKNOWLEDGMENTS

We would like to thank J. Benichov and N. Hein for their help with fieldwork; T. Eliav and A. Navas for providing helpful comments to the project and manuscript; and A. Costalunga for the drawings of nightingales.

Funding sources: The Joachim Herz Stiftung Add-on Fellowships for Interdisciplinary Life Science - awarded to G.C.; The ERC Consolidator Grant 819603 SYNAPSEEK - awarded to T.P.V.; The HORIZON EUROPE European Research Council (ERC)-2017-StG-757459 MIDNIGHT, Deutsche Forschungsgemeinschaft VA742/2-1, Deutsche Forschungsgemeinschaft 327654276–SFB 1315 - awarded to D.V.

## AUTHOR INFORMATION

These authors contributed equally: J.S. Calderon-Garcia & G. Costalunga

### Authors and Affiliations

Institute of Science and Technology Austria, Klosterneuburg 3400, Austria Calderon-Garcia J.S. & Vogels T.P.

Neural Circuits for Vocal Communication Research Group, Max Planck Institute for Biological Intelligence, Eberhard-Gwinner-Str., Seewiesen 82319, Germany Costalunga G. & Vallentin D.

### Contributions

J.S.C.G., G.C., and D.V. conceived, designed, conducted the experiments, analysed the data and crafted the figures; J.S.C.G. and G.C. wrote the first draft of the manuscript. All authors participated in writing and editing the manuscript. G.C., T.P.V. and D.V. acquired funding; T.P.V. and D.V. supervised the project.

### Corresponding author

Correspondence to Daniela Vallentin

## DECLARATION OF INTERESTS

The authors declare no competing interests.

## SUPPLEMENTARY INFORMATION

**Figure S1.**
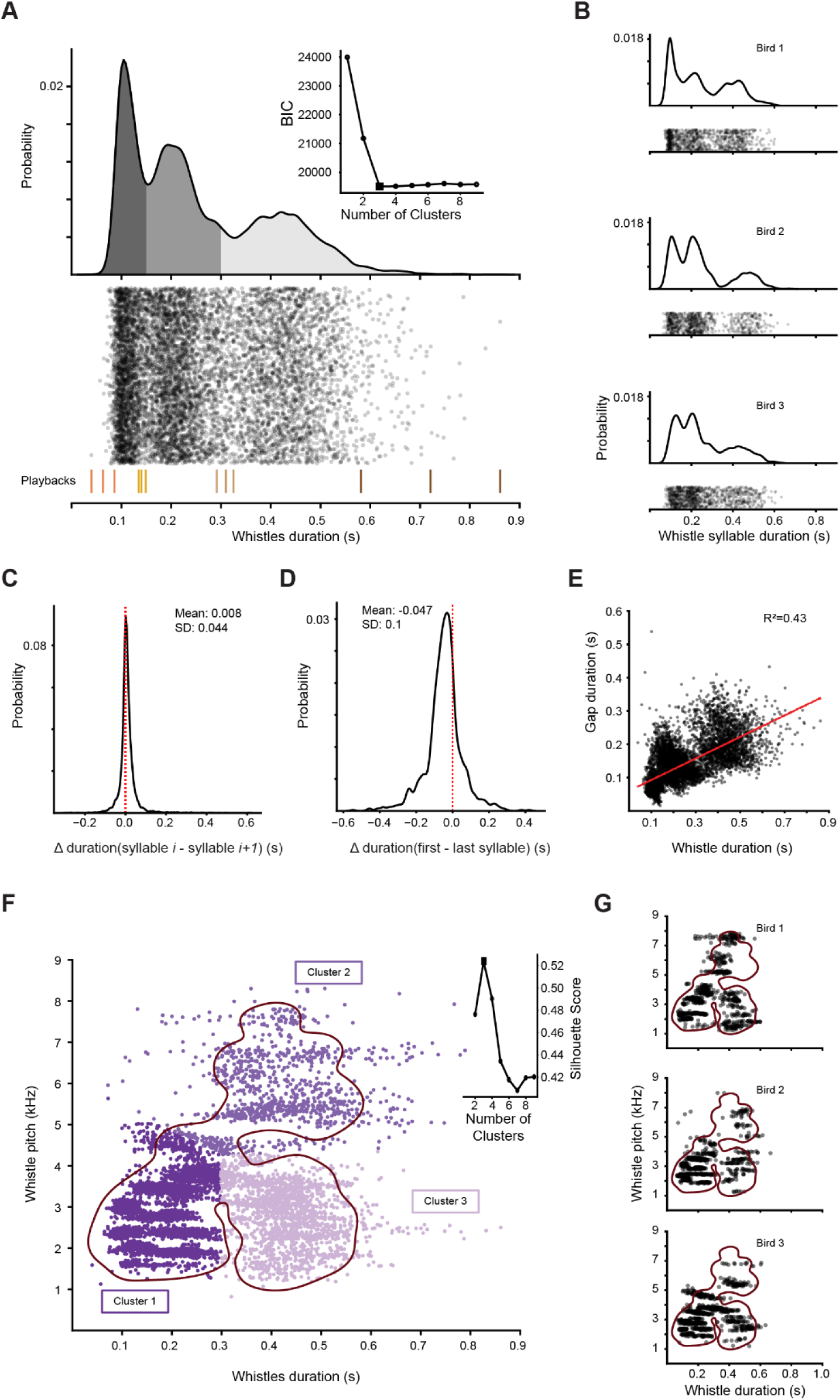
Temporal features of whistle songs in naturally singing nightingales. (A) Natural distribution of duration of whistle syllables (0.217±0.098 s, n=8448 syllables from N=11 birds). The three grey colours represent the three modes of the probability distribution defined by a Gaussian Mixture Model (See methods). Each datapoint corresponds to the duration of a single whistle syllable. Vertical colour lines represent the duration of the playbacks for the first experiment. The inset plot shows the Bayesian Information Criterion to define the optimal number of modes. (B) Natural distribution of duration of whistle syllables for three different example birds: Bird 1 (0.220±0.118 s, n=1041), Bird 2 (0.208±0.077 s, n=875), and Bird 3 (0.213±0.079 s, n=1117). Each datapoint corresponds to the duration of a single whistle syllable. (C) Natural distribution of the difference in duration between subsequent whistles in a song (0.004±0.012 s). The red dashed line outlines the zero difference. (D) Natural distribution of the difference in duration between the first and the last whistle in a whistle song (-0.041±0.04 s). The red dashed line outlines the zero difference. (E) Relationship between whistle duration and the duration of the subsequent silent interval from the control data (n=7209). Each datapoint represents an individual whistle syllable and the respective following silent gap. The red line represents the linear relationship. (F) Natural distribution of whistle syllable duration and pitch. Each datapoint represents the duration and pitch of a single whistle syllable (n=8448, N=11). The three colours represent the clusters defined by a k-means algorithm (See methods, Cluster 1 centers at 0.168 s/2.799 kHz; Cluster 2 centers at 0.412 s/5.817 kHz; Cluster 3 centers at 0.431 s/2.819 kHz). The brown line encloses 90% of the control KDE density shown in Fig 1E. The inset plot shows the Silhouette score to define the optimal number of clusters. (G) Natural distribution of whistle syllable duration and pitch for three different example birds. Each datapoint corresponds to the duration and pitch of a single whistle syllable (n=1041, 875 and 1117). The brown line encloses 90% of the control KDE density shown in Fig 1E.

**Figure S2.**
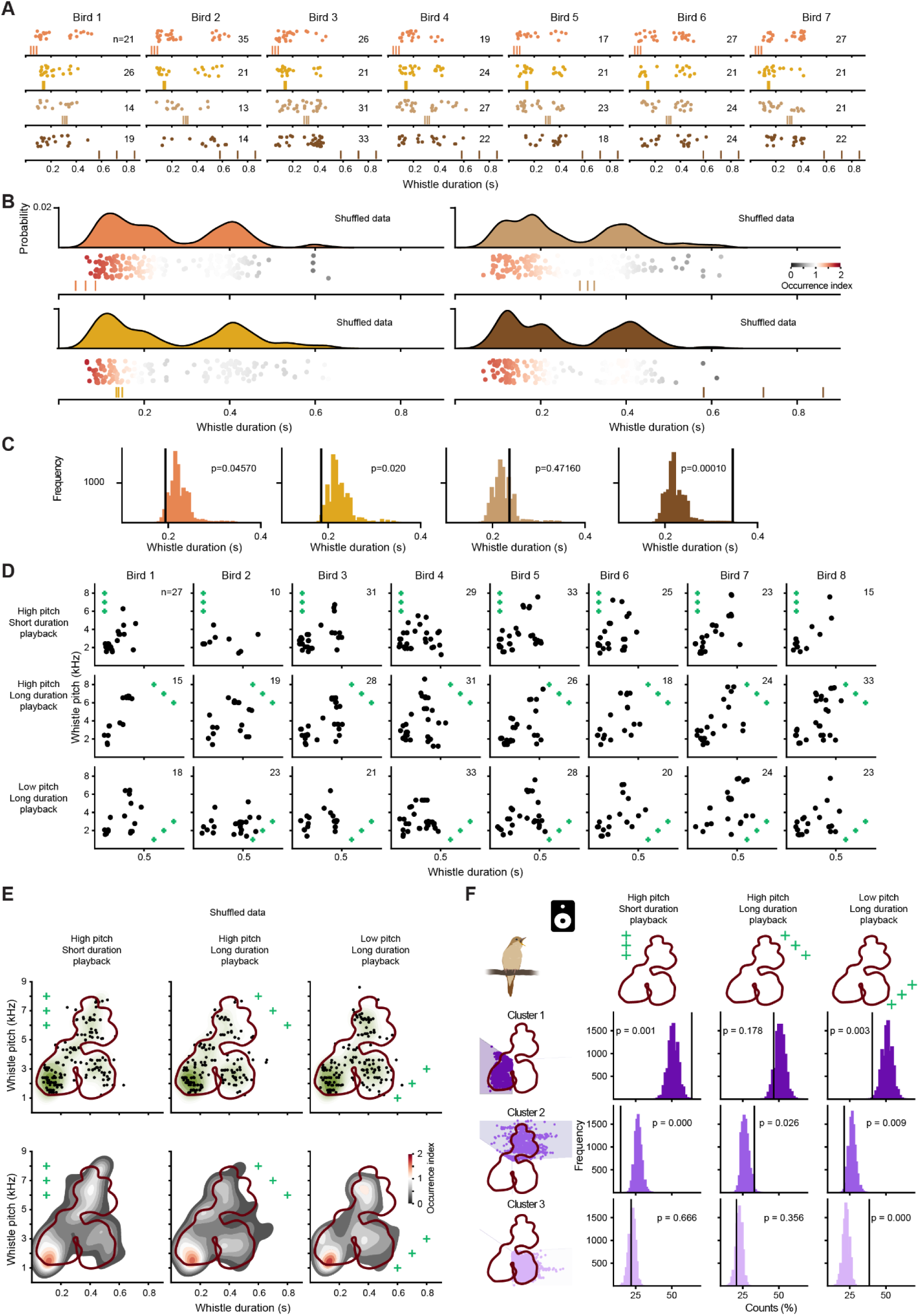
Experimental data and statistical analysis on imitation of whistle syllable duration. (A) Distribution of duration of whistle responses to each playback regime for all birds: Very-Short (Bird 1, 0.179±0.087 s; Bird 2, 0.2±0.096 s; Bird 3, 0.165±0.054 s; Bird 4, 0.194±0.054 s; Bird 5, 0.153±0.081 s; Bird 6, 0.345±0.127 s; Bird 7, 0.327±0.104 s); Medium-Short (0.18±0.040 s; 0.191±0.107 s; 0.15± 0.033 s; 0.220± 0.119 s; 0.185± 0.104 s; 0.368±0.119 s; 0.162± 0.041 s); Medium-Long (0.255±0.093 s; 0.235±0.055 s; 0.275±0.118 s; 0.210±0.121 s; 0.232±0.120 s; 0.367±0.115 s; 0.188±0.69 s), Very-Long (0.194±0.07 s; 0.489±0.071 s; 0.377±0.045 s; 0.271±0.133 s; 0.365±0.049 s; 0.412±0.094 s; 0.296±0.087 s). Each datapoint represents the median whistle syllable duration of whistle syllables within a song in response to a playback. Data points are color-coded based on the playback regime. (B) Example of probability distributions of shuffled responses to each playback regime: Very-Short (0.214±0.112 s); Medium-Short (0.221±0.102 s); Medium-Large (0.214±0.122 s); Very-Large (0.218±0.110 s). Each data point represents the median whistle syllable duration of whistle syllables within a song in response to a playback. Datapoints are color-coded based on occurrence index of whistle syllable durations (See methods). (C) Distribution of medians, calculated from 10000 permutations of shuffled data for each playback regime. Observed medians are indicated with black lines. (D) Scatter plot of whistle syllable duration and pitch in response to the three playback types for all the birds. Each datapoint represents the median duration and pitch of whistle syllables within a song. The green crosses represent the duration and pitch of the three types of playbacks (High-pitch/Short-duration, High-pitch/Long-duration, Low-pitch/Long-duration). (E) Top: Example distribution of whistle syllable duration and pitch of shuffled responses in response to the three playback types, represented as in Fig 1E. Bottom: Density estimation of shuffled responses color-coded based on the relative density of occurrences compared to the naturally observed distribution, calculated as the occurrence index of observed to expected densities (See methods). The brown line encloses 90% of the control KDE density. (F) Distribution of % of counts on each cluster in response to each playback type, calculated from 10000 permutations of shuffled data. Observed % of counts are indicated with the black lines (High-pitch/Short-duration: C1 63.7%, C2 14.5%, C3 21.8%; High-pitch/Long-duration: C1 46.4%, C2 33%, C3 20.6%; Low-pitch/Long-duration C1 40.5%, C2 21.1%, C3 38.4%).

## Notes

### Competing Interest Statement

The authors have declared no competing interest.

